# Adaptive Laboratory Evolution (ALE) enables carbon-negative mixotrophic fermentation and enhanced chain elongation in *Clostridium* sp. JS66

**DOI:** 10.64898/2026.08.16.744336

**Authors:** Ji Myeong Kim, Taekwan Moon, Jung Ho Ahn, Ja Kyong Ko, Gyeongtaek Gong, Jae Yong Ryu, Sung Ok Han, Min-Kyu Oh, Youngsoon Um

## Abstract

Improving carbon recovery during sugar fermentation remains a major challenge because a substantial fraction of substrate carbon is lost as CO_2_ during central metabolism. To overcome this limitation, *Clostridium* sp. JS66 (JS66), an acetogen producing hexanoic acid from glucose, was subjected to adaptive laboratory evolution under CO_2_/H_2_ conditions to enhance H_2_-assisted CO_2_ reassimilation during glucose fermentation. The evolved strain, ALECO2, exhibited CO_2_ consumption without a lag phase under autotrophic conditions and reached a 9.5-fold higher CO_2_ uptake rate than JS66. Under fed-batch conditions, glucose-only fermentation yielded a carbon molar yield (C_metabolite_/C_sugar,_ C_M_/C_S_) of 0.60, whereas H_2_ supplementation increased C_M_/C_S_ to 0.91 and redirected carbon flux toward C6 products (hexanoic acid and hexanol), which accounted for 49% of total C_output. With additional CO_2_ supplementation, ALECO2 further assimilated externally supplied CO_2_, increasing the C_M_/C_S_ to 1.10 and demonstrating carbon-negative fermentation. Assimilation of externally supplied CO_2_ further redirected carbon flux toward chain elongation, producing 7.14 g/L hexanoic acid and increasing the C6 carbon fraction to 57% of total C_output. Constraint-based flux analysis supported increased acetyl-CoA formation through the Wood-Ljungdahl pathway and enhanced flux toward reverse β-oxidation under H_2_- and CO_2_/H_2_-supplemented conditions. Genome analysis identified mutations including genes encoding a putative HytB homolog and a LysR-type transcriptional regulator. These results establish ALECO2 as a promising evolved anaerobic non-photosynthetic (ANP) mixotrophy platform that links CO_2_ reassimilation and external CO_2_ assimilation with chain elongation, enabling carbon-neutral and carbon-negative production of value-added C6 products from glucose.

## 1. Introduction

Global CO_2_ emissions reached a record 37.2 Gt in 2025, highlighting the need for sustainable bioprocesses that produce chemicals from renewable feedstocks while reducing carbon emissions (Deng et al., 2026). Lignocellulosic sugars are attractive feedstocks because they are abundant, renewable, and do not compete with food resources (Wijaya et al., 2014). However, conventional sugar fermentation is intrinsically constrained by carbon loss. During glycolysis, glucose-derived pyruvate is converted to acetyl-CoA with concomitant CO_2_ release, and one-third of the substrate carbon can be lost. As a result, the theoretical carbon recovery into acetyl-CoA-derived products is limited to approximately 67% in conventional heterotrophic fermentation (Jones et al., 2016). Thus improving carbon recovery through reassimilation of glycolytically produced CO_2_ while retaining the practical advantages of sugar fermentation represents an attractive strategy for developing carbon-conserving bioprocesses.

Acetogenic bacteria offer a promising approach for overcoming this limitation because they naturally possess the Wood-Ljungdahl pathway (WLP), which converts CO_2_ into acetyl-CoA. Several strategies have been developed to improve carbon recovery during sugar fermentation by exploiting acetogenic CO_2_ reassimilation. One strategy is the use of synthetic co-culture platforms that combine sugar-fermenting *Clostridium* strains with CO_2_/H_2_-utilizing acetogens. In these systems, the sugar-fermenting partner converts carbohydrates into acids and alcohols while releasing CO_2_ and H_2_, whereas the acetogenic partner reassimilates these gases through the WLP. Such division of metabolic labor has been shown to improve carbon recovery, expand product spectra, and, in some cases, achieve carbon recoveries exceeding 100% through the incorporation of externally supplied CO_2_ and H_2_ (Charubin et al., 2019; Li et al., 2024; Willis et al., 2026). Nevertheless, co-culture platforms introduce additional operational complexity because their performance depends on coordinating physiologically distinct organisms. Parameters such as population ratio, inoculation timing, growth-rate balance, and interspecies metabolite exchange can strongly influence process outcome, which may complicate process control and reproducibility (Song et al., 2024).

Alternatively, anaerobic non-photosynthetic (ANP) mixotrophy performed by acetogenic monocultures provides a simpler strategy for improving carbon recovery through the simultaneous utilization of sugars and inorganic substrates (i.e., CO_2_/H_2_, CO) (Fast et al., 2015). During ANP mixotrophy, endogenously produced CO_2_ can be reassimilated into acetyl-CoA through the WLP using reducing equivalents derived from H_2_ generated during sugar catabolism, allowing carbon recovery into metabolites to exceed the theoretical maximum of 67% expected from heterotrophic fermentation (Jones et al., 2016). However, the reducing equivalents generated during sugar fermentation are generally insufficient to support complete reassimilation of glucose-derived CO_2_ through the WLP. Consequently, exogenous H_2_ has been employed as an electron donor to provide the additional reducing power required for complete CO_2_ fixation and improved carbon conservation. Several studies have demonstrated H_2_-assisted CO_2_ reassimilation in acetogenic monocultures, including *Clostridium ljungdahlii* and *Blautia producta*, achieving carbon recoveries approaching 100% (Jones et al., 2016; Maru et al., 2018). Despite these advances, these systems primarily produce short-chain C2-C3 compounds, limiting the application of ANP mixotrophy to higher-value products. Expanding ANP mixotrophy toward chain-elongated products would considerably broaden its industrial applicability. Among these products, hexanoic acid (HA), a medium-chain carboxylic acid, is particularly attractive because of its higher energy density than short-chain acids and its applications as a food flavoring agent, fragrance ingredient, and precursor for hexyl esters and hexanol (HeOH), which are used as fuel additives and chemical intermediates (Kim et al., 2022). Furthermore, the model acetogens commonly used for ANP studies, such as *C. ljungdahlii* and *C. autoethanogenum*, utilize fructose rather than glucose as their fermentable sugar (Jones et al., 2016; Maru et al., 2018). This highlights the need to extend ANP mixotrophy toward chain elongation in glucose-utilizing acetogens for the conversion of glucose-rich lignocellulosic biomass into value-added medium-chain products. To date, no acetogenic monoculture has been reported to simultaneously achieve H_2_-assisted CO_2_ reassimilation and enhanced production of medium-chain compounds from glucose.

This study aimed to expand ANP mixotrophy toward HA production using *Clostridium* sp. JS66 (JS66), an acetogen that produces HA as the main product from glucose, as a monoculture platform (Kim et al., 2022). JS66 was subjected to adaptive laboratory evolution (ALE) under CO_2_/H_2_ conditions to enhance gas utilization, yielding the evolved strain ALECO2. The ANP mixotrophy performance of ALECO2 was evaluated under glucose-fed, glucose/H_2_-fed, and glucose/CO_2_/H_2_-fed mixotrophic conditions to assess carbon conservation, the feasibility of carbon-negative fermentation, and carbon flux redistribution. Finally, metabolic flux analysis and whole-genome sequencing of ALECO2 were conducted to identify the mechanisms underlying enhanced chain elongation and gas utilization. This work provides a monoculture strategy for coupling CO_2_ reassimilation with chain elongation and demonstrates the potential of ANP mixotrophy for carbon-conserving and carbon-negative production of HA from glucose.

## 2. Materials and methods

### 2.1. Strains, media, and cultivation

*Clostridium* sp. JS66 was cultivated following a previously described method (Kim et al., 2022). The parental strain used for adaptation was ALECO70, an evolved variant of JS66 obtained through adaptive laboratory evolution under high CO conditions containing 70% CO. Glycerol stocks were stored at -80 ℃ and inoculated at 5% (v/v) into 57 mL serum bottles containing 20 mL of modified P2 medium for seed culture with an initial pH of 6.0 adjusted using 2 M KOH, as previously described (Kim et al., 2022). Seed cultures were incubated at 30 ℃ with shaking at 150 rpm.

Modified P7 medium (MP7-S) was used for main cultures. The medium was prepared according to Oh et al. (2026), except that CaCO_3_ was added to enhance pH-buffering capacity. Main cultures were inoculated at an initial optical density at 600 nm (OD_600_) of 0.2. For strain screening after ALE, single colonies were isolated using 2X YTG agar plates composed of 5 g/L glucose, 16 g/L tryptone, 10 g/L yeast extract, 0.9 g/L NaCl, 0.001 g/L resazurin, and 17 g/L agarose.

Fermentations were performed in 157 mL serum bottles containing 20 mL of MP7-S medium with 2 g/L CaCO_3_ for batch fermentation or 4 g/L for fed-batch fermentation. Cultures were incubated at 30 ℃ with shaking at 200 rpm. In glucose-fed mixotrophic fermentation, glucose was supplied at 3 g/L every 24 h without external gas supplementation. In glucose/H_2_-fed mixotrophic fermentation, glucose was supplied at 3 g/L every 24 h, whereas H_2_ was initially supplied to 150 kPa and replenished to 150 kPa every 24 h when the headspace H_2_ pressure decreased. In glucose/CO_2_/H_2_-fed mixotrophic fermentation, glucose (3 g/L) and H_2_ (150 kPa) were supplied every 24 h, while CO_2_ was initially adjusted to a partial pressure of 20 kPa and replenished to 20 kPa when the headspace CO_2_ pressure decreased.

For *in situ* product adsorption experiments, GSP-25 resin was added at 200 g/L. Resin pretreatment and preparation of resin-containing MP7-S medium were performed according to Oh et al. (2026). After fermentation, the culture supernatant was removed, and resin-adsorbed products were recovered by extraction with 20 mL of ethyl acetate at 30 ℃ and 150 rpm for 1 h. Since the broth and ethyl acetate phases were at equal volumes, the total titer of each metabolite (acetic acid (AA), butyric acid (BA), HA, ethanol (EtOH), butanol (BuOH), and HeOH) was determined as the sum of its concentrations in the two phases, following the previously reported method (Jung et al., 2026).

### 2.2. Adaptive laboratory evolution to enhance gas utilization

ALE for enhanced CO_2_ consumption was performed using ALECO70 as the parental strain. Cultures were grown in MP7-S medium under a pressurized gas mixture of CO_2_ and H_2_ (30:70). Serial transfers were carried out when more than 50% of the supplied CO_2_ had been consumed. To progressively increase selection pressure for gas utilization, cultivation parameters including gas pressure, transfer volume, bottle volume, and CaCO_3_ concentration were systematically adjusted throughout the ALE process (Supplementary material-Excel file). All cultures were incubated at 30 ℃ with shaking at 200 rpm. After 95 rounds, ten individual colonies were randomly isolated and screened for CO_2_ utilization under autotrophic conditions (CO_2_:H_2_ = 30:70, 200 kPa). The colony exhibiting the fastest complete consumption of CO_2_ was selected and designated ALECO2.

### 2.3. Genome sequencing

To identify genetic mutations accumulated during ALE, whole-genome resequencing (WGrS) of ALECO70 and ALECO2 was performed using the Illumina HiSeq X Ten platform. The resulting 151-bp paired-end reads were aligned to the *Clostridium* sp. JS66 reference genome (RefSeq assembly NZ_CP131467.1), and variants were identified using Breseq v0.36.0 (Deatherage et al., 2014). For *de novo* genome assembly, PacBio Revio HiFi reads were generated from high-molecular-weight genomic DNA. ALECO70 reads were assembled using Flye v2.9, whereas ALECO2 reads were assembled using the Microbial Genome Analysis workflow in SMRT Link v25.1 with default parameters (Chin et al., 2013; Kolmogorov et al., 2019).

### 2.4. Flux balance analysis

Flux balance analysis (FBA) was performed using a previously reported core metabolic model of *C. carboxidivorans* P7 (Vees et al., 2022). Experimentally measured glucose uptake rates, H_2_ and CO_2_ exchange rates, and AA and EtOH production rates were used as condition-specific constraints in mmol/gDCW/h, where gDCW denotes grams of dry cell weight. Flux sampling was conducted using the optGpSampler to explore feasible intracellular flux distributions under each condition, with 1500 samples generated per condition (Megchelenbrink et al., 2014). All simulations were performed using the COBRApy python package ver. 0.30.0 (Ebrahim et al., 2013).

### 2.5. Analytical methods and calculations

Analytical methods were performed as described previously, with minor modifications (Jung et al., 2026; Oh et al., 2026). Cell growth was monitored by OD_600_ measurement using a UV-visible spectrophotometer. When OD_600_ measurement was hindered by CaCO_3_ supplementation, cell growth was estimated from total protein concentration. Cells were lysed using a freeze-thaw procedure modified from Fagan et al. (2011), and soluble protein was quantified using a bicinchoninic acid assay. OD_600_ values were then estimated using a pre-established calibration curve between OD_600_ and total protein concentration. Gas consumption rates were calculated from changes in headspace partial pressure using the ideal gas law and expressed as mmol/h. Carbon proportion (%) was calculated as the carbon moles recovered in C2 (AA and EtOH), C4 (BA and BuOH), or C6 (HA and HeOH) compounds divided by the total carbon moles recovered in all C2-C6 products. Carbon molar yield (C_metabolite_/C_sugar,_ C_M_/C_S_) was calculated by dividing the total carbon moles recovered in C2-C6 acids/alcohols by the carbon moles of glucose consumed. In fed-batch fermentations, C_input represents the cumulative carbon consumed from glucose and, where applicable, the net carbon consumed from externally supplied CO_2_ based on the decrease in headspace CO_2_. C_output represents the cumulative carbon recovered in fermentation products, cell biomass, and net CO_2_ produced in the headspace.

### 2.6. Statistical analysis

Statistical analyses of experimental data were performed in R version 4.5.2 using Welch’s t-test for pairwise comparisons and one-way analysis of variance (ANOVA) followed by Tukey’s post hoc test for multiple-group comparisons. Flux distributions obtained from random sampling analysis were compared using the Kruskal-Wallis test. Significance was defined as *p* < 0.05.

## 3. Results and discussion

### 3.1. Mixotrophic behavior of *Clostridium* sp. JS66

In acetogens, sugar catabolism and reassimilation of sugar-derived CO_2_ can occur simultaneously (ANP mixotrophic metabolism). As a result, C_M_/C_S_ can exceed the theoretical value of 0.67 expected from heterotrophic fermentation, although the extent of CO_2_ reassimilation varies depending on the strain and cultivation conditions (Jones et al., 2016; Maru et al., 2018). To characterize the mixotrophic behavior of JS66, batch fermentation was performed in serum bottles containing 5 g/L glucose, and changes in headspace gas composition and metabolites were monitored. Glucose was completely consumed within 24 h and C2-C6 acids/alcohols were produced (Supplementary material-Word file). During the initial 24 h of fermentation, both CO_2_ and H_2_ accumulated in the headspace concomitantly with glucose consumption. After 48 h, the pressure of CO_2_ and H_2_ gradually decreased, accompanied by a continuous increase in AA concentration (Supplementary material-Word file), suggesting utilization of CO_2_ and H_2_ via WLP. The C_M_/C_S_ value at 24 h was 0.59, suggesting that CO_2_ reassimilation during the glucose consumption phase was limited and that simultaneous utilization of glucose and gaseous substrates was not substantial. One possible explanation for this apparent limitation of mixotrophic metabolism is that JS66 preferentially utilized glucose while WLP-mediated CO_2_ fixation remained relatively weak during 24 h of fermentation. Carbon catabolite repression (CCR) has been reported in some acetogens, such as *C. aceticum*, in which only limited CO_2_ consumption occurred in the presence of fructose (Braun et al., 1981). In addition, the extent of mixotrophic metabolism is known to depend not only on the microbial strain but also on cultivation conditions. For example, *B. producta* exhibited a C_M_/C_S_ value of 0.63 in serum bottle cultivation with glucose 10 g/L, whereas a substantially higher C_M_/C_S_ value of 0.90 was achieved in a pH-controlled bioreactor supplied with 20 g/L glucose (Maru et al., 2018).

Alternatively, the observed accumulation of CO_2_ during glucose fermentation may reflect an intrinsically low rate of gas utilization relative to the rapid generation of CO_2_ from glucose metabolism (Supplementary material-Word file). To evaluate this possibility, the gas consumption behavior of JS66 was examined under autotrophic conditions in the absence of glucose. JS66 consumed CO_2_ and H_2_ after an initial lag period, with approximately 10 kPa CO_2_ consumed between 48 and 96 h (Supplementary material-Word file), whereas approximately 20 kPa CO_2_ had accumulated in the headspace by 24 h of glucose fermentation (Supplementary material-Word file). These results suggest that the intrinsic rate of CO_2_ utilization by JS66 was insufficient to offset the rapid generation of CO_2_ during glucose fermentation, thereby limiting apparent CO_2_ reassimilation during the glucose consumption phase. Collectively, these observations suggested that enhancing the CO_2_ and H_2_ consumption rate could promote more effective co-utilization of glucose and gas substrates.

### 3.2. Adaptive laboratory evolution for enhancing CO_2_ consumption rate of JS66

Based on these observations, ALE was employed to improve CO_2_ and H_2_ utilization by JS66 and thereby enhance mixotrophic carbon fixation. Prior to ALE, the CO_2_ and H_2_ utilization capacities of the wild-type JS66 strain and ALECO70 were compared under a CO_2_:H_2_ gas mixture (30:70, 150 kPa). ALECO70 had previously been evolved under 70% CO and exhibited an increase in CO consumption rate from 0.18 to 0.52 mmol CO/h. Both strains exhibited a lag phase of approximately 48 h (Supplementary material-Word file). Following this lag phase, however, ALECO70 showed substantially higher gas consumption than JS66, consuming 27.1 kPa of CO_2_ compared to 10.5 kPa for JS66 by 96 h (Supplementary material-Word file). The enhanced utilization of CO_2_ and H_2_ by ALECO70 suggests that adaptation to CO might improve the overall capacity for gas metabolism. Therefore, ALECO70 was selected as the parental strain for subsequent evolution aimed at improving CO_2_ and H_2_ utilization.

During ALE under a CO_2_/H_2_ gas mixture (30:70) (Supplementary material-Excel file), the evolved population exhibited progressive improvements in gaseous substrate utilization. The CO_2_ consumption rate increased from 0.021 mmol CO_2_/h in the initial ALE round to 0.21 mmol CO_2_/h by the 95th round, representing a 10-fold improvement (Fig. 1A). Similarly, the H_2_ consumption rate increased from 0.05 to 0.56 mmol H_2_/h during the same period (Fig. 1B). Following completion of ALE, ten colonies were randomly isolated from the 95th round and screened for CO_2_ utilization. The colony exhibiting the highest CO_2_ uptake under autotrophic conditions was selected and designated ALECO2.

**Figure 1.**
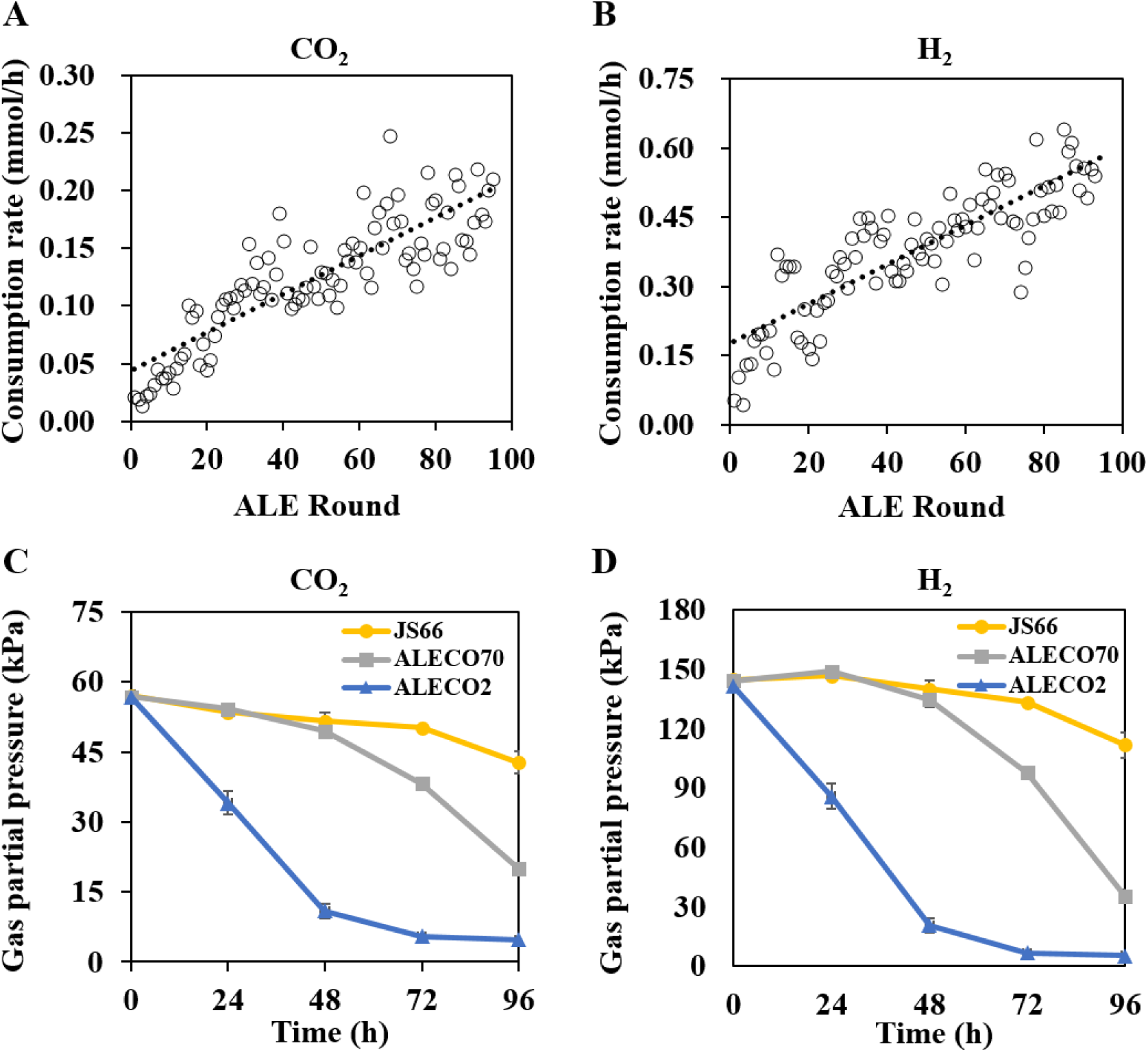
CO_2_ and H_2_ consumption dynamics during and after adaptive laboratory evolution (ALE). (A) Specific CO_2_ and (B) H_2_ consumption rates during the exponential phase across successive ALE rounds. Each dot represents a single ALE round, and dotted lines show overall trends of increased gas uptake capacity. (C) CO_2_ and (D) H_2_ consumption profiles during autotrophic fermentation by the *Clostridium* sp. JS66, ALECO70, and ALECO2 strains. Error bars represent the range of two biological replicates (n = 2).

To verify the enhanced gas-utilization phenotype of the ALECO2 strain, batch gas fermentation was performed using JS66, ALECO70, and ALECO2 with a CO_2_:H_2_ gas mixture (30:70, 200 kPa). Compared with the JS66 and ALECO70, ALECO2 exhibited markedly improved gaseous substrate utilization, consuming both CO_2_ and H_2_ from the beginning of cultivation (Fig. 1C-D). In contrast, JS66 and ALECO70 displayed delayed gas consumption, with uptake occurring after 48-72 h of cultivation. The ALECO2 strain exhibited a CO_2_ consumption rate of 0.053 mmol CO_2_/h during 24-48 h, which was 4.8-fold and 9.5-fold higher than those of ALECO70 (0.011 mmol CO_2_/h) and JS66 (0.0056 mmol CO_2_/h), respectively. Consistent with the enhanced CO_2_ consumption, H_2_ uptake was also substantially accelerated in ALECO2 (Fig. 1D). These results demonstrate that ALE effectively enhanced the capacity of JS66 for CO_2_ and H_2_ utilization.

To evaluate whether enhanced gas utilization translated into improved mixotrophic metabolism, ALECO2 was cultivated in serum bottles containing 5 g/L glucose. Similar to JS66, glucose was completely consumed within 24 h (Supplementary material-Word file). Unlike JS66, however, ALECO2 achieved a C_M_/C_S_ of 0.71 at 24 h, exceeding the theoretical heterotrophic value of 0.67. This result indicates that reassimilation of glucose-derived CO_2_ occurred concurrently with glucose metabolism, although net CO_2_ accumulation in the headspace indicates that CO_2_ production exceeded its reassimilation during the initial stage of fermentation. After glucose depletion, the partial pressures of CO_2_ and H_2_ gradually decreased after 48 h, indicating the occurrence of gaseous substrate utilization during the later stage of fermentation. By 96 h, H_2_ was completely consumed, whereas approximately 16 kPa CO_2_ remained in the headspace at the same time point (Supplementary material-Word file), suggesting that H_2_ availability became the limiting factor for further CO_2_ reassimilation. Collectively, these results indicate that ALE enhanced the capacity for gaseous substrate utilization and promoted more efficient mixotrophic metabolism in ALECO2.

### 3.3. Carbon-neutral fermentation by H_2_-mixotrophy using ALECO2 strain (batch fermentation)

As demonstrated in Section 3.2, ALECO2 completely consumed H_2_ while residual CO_2_ remained in the headspace (Supplementary material-Word file), suggesting that the availability of reducing equivalents may have limited further CO_2_ reassimilation. To evaluate whether exogenous H_2_ supplementation could overcome this limitation and promote more complete carbon conservation, batch H_2_-mixotrophic fermentation was performed with JS66, ALECO70, and ALECO2 under identical conditions. Glucose (5 g/L) was supplied as the carbon source, and H_2_ was provided as an external electron donor to facilitate CO_2_ fixation via the WLP.

All strains completely consumed glucose within 24 h, accompanied by CO_2_ production (Fig. 2A-C). However, their gas profiles differed markedly among the strains. At 24 h, JS66 and ALECO70 had accumulated 16-17 kPa CO_2_, whereas ALECO2 accumulated only 3.5 kPa (Fig. 2A-C), suggesting that enhanced WLP activity in ALECO2 partially offset CO_2_ generation even during active glycolysis. By 48 h, CO_2_ was completely reassimilated in ALECO2 (Fig. 2C), while residual CO_2_ remained in JS66 and ALECO70 throughout the fermentation, reaching final partial pressures of 15.8 and 6.3 kPa at 96 h, respectively (Fig. 2A-B). Concurrently, the H_2_ partial pressure decreased by 53.5 kPa in ALECO2 by 96 h, compared with decreases of only 0.2 and 24.5 kPa in JS66 and ALECO70, respectively (Fig. 2A-C). Notably, the substantially lower CO_2_ accumulation in ALECO2 at 24 h under H_2_-mixotrophic conditions (3.5 kPa) compared with glucose-only fermentation (∼19 kPa, Supplementary material-Word file) confirms that H_2_ availability was the primary constraint on CO_2_ reassimilation in the absence of external H_2_ supply.

**Figure 2.**
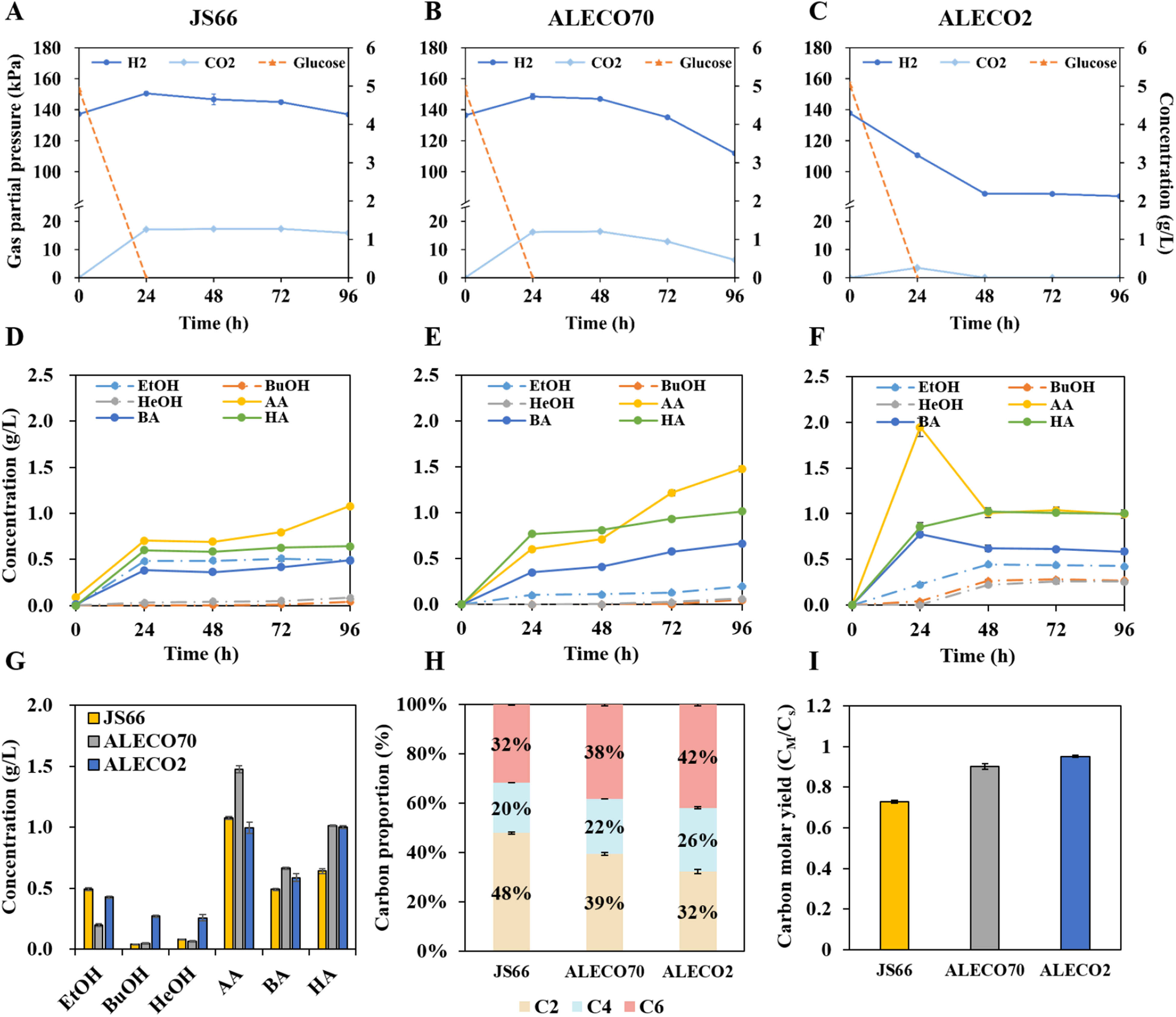
H_2_-mixotrophic batch fermentation performance of *Clostridium* sp. JS66, ALECO70, and ALECO2 strains. (A-C) Time-course profiles of CO_2_ and H_2_ partial pressures and glucose concentrations of (A) JS66, (B) ALECO70, and (C) ALECO2. (D-F) Time-course profiles of ethanol (EtOH), butanol (BuOH), hexanol (HeOH), acetic acid (AA), butyric acid (BA), and hexanoic acid (HA) produced by each strain. (G) Final concentrations of fermentation products. (H) Carbon proportion (%) of C2 (AA and EtOH), C4 (BA and BuOH), and C6 (HA and HeOH) compounds based on the total carbon recovered in all C2-C6 products. (I) Carbon molar yields (C_M_/C_S_), defined as the ratio of carbon moles in produced metabolites to those in consumed glucose, for each strain. Error bars represent the range of two biological replicates (n = 2).

Time-course product analysis revealed distinct carbon redistribution patterns among the three strains. In JS66 and ALECO70, C2-C6 acids/alcohols were produced, and AA continued to accumulate after glucose depletion concomitantly with CO_2_ and H_2_ consumption, reaching final concentrations of 1.08 and 1.48 g/L in JS66 and ALECO70, respectively (Fig. 2D, 2E, and 2G). In contrast, ALECO2 produced AA and BA during the initial 24 h of glucose fermentation, but both acids subsequently decreased between 24 and 48 h, coinciding with complete CO_2_ reassimilation and substantial H_2_ consumption (Fig. 2C and 2F). During the same period (24-48 h), EtOH and BuOH increased as AA and BA declined, suggesting that part of the accumulated AA and BA was reassimilated and subsequently redirected toward alcohol formation. In contrast, HA and HeOH accumulated simultaneously, suggesting that chain elongation toward C6 products continued actively during this period, resulting in expansion of the C6 product pool rather than simple conversion of HA to HeOH (Fig. 2F).

Analysis of the product carbon proportion showed that C2 compounds remained the major products in JS66 and ALECO70, accounting for 48% and 39% at 96 h, respectively, (Fig. 2H). In comparison, JS66 and ALECO70 exhibited lower carbon proportion of C6 products (32% and 38%, respectively) than ALECO2 (42%) (Fig. 2H). Together, these observations suggest that the elevated H_2_ consumption in ALECO2 supported not only CO_2_ reassimilation through the WLP but also the reductive reactions required for chain elongation toward C6 products and alcohol formation during the period of active CO_2_ reassimilation following glucose depletion.

The C_M_/C_S_ value further demonstrated the superior carbon conservation achieved by ALECO2. The JS66 exhibited a C_M_/C_S_ value of 0.72, slightly higher than the theoretical value of 0.67 expected from glucose catabolism without CO_2_ reassimilation (Fig. 2I). The ALECO70 strain exhibited a higher C_M_/C_S_ value of 0.90, suggesting substantial but incomplete CO_2_ reassimilation. Notably, the ALECO2 strain achieved a C_M_/C_S_ value of 0.95 under the H_2_-mixotrophic conditions (Fig. 2I). This value approaches the theoretical maximum of 1.0, indicating near-complete recovery of glucose-derived carbon through reassimilation of fermentation-derived CO_2_. Considering that a portion of glucose-derived carbon was incorporated into biomass and minor metabolites that were not quantified, the observed C_M_/C_S_ value of 0.95 represents near-complete carbon recovery. Furthermore, no net accumulation of CO_2_ was detected at the end of fermentation (Fig. 2C), demonstrating that exogenous H_2_ enabled effective reassimilation of metabolically generated CO_2_.

Several studies have explored H_2_-mixotrophic fermentation in which exogenous H_2_ serves as an electron donor for CO_2_ reassimilation through the WLP. For example, the engineered acetone-producing strain *C. ljungdahlii* ΔSADH (pTCtA) achieved carbon molar yields approaching 100% when cultivated with fructose and 40% of H_2_ in the headspace of serum bottles, indicating near-complete reassimilation of fermentation-derived CO_2_ (Jones et al., 2016). Similarly, *B. producta* achieved a carbon molar yield of 98.3% during H_2_-mixotrophic fermentation with 20 g/L glucose in a 3-L bioreactor supplied with H_2_ (Maru et al., 2018). In contrast, *C. carboxidivorans*, a producer of C2-C6 acids and alcohols, achieved a carbon yield of only 83.4% during continuous H_2_-mixotrophic fermentation with glucose in a 2-L bioreactor, indicating that a considerable fraction of carbon was still lost as CO_2_ (Vees et al., 2022).

Although direct comparison among these studies is difficult because of differences in strain background and cultivation conditions, several notable distinctions can be identified. The H_2_-mixotrophic systems based on *C. ljungdahlii* and *B. producta* primarily produced C2-C3 compounds, whereas ALECO2 maintained active chain elongation toward C6 products while simultaneously achieving near-complete carbon recovery. Vees et al. (2022) reported that C2 compounds (AA and EtOH) were the main products in H_2_-mixotrophic fermentation of *C. carboxidivorans*. In contrast, ALECO2 exhibited a shift in carbon flux toward C6 products, increasing the carbon proportion of C6 compounds to 42% while achieving a C_M_/C_S_ value of 0.95. These results suggest that ALECO2 not only effectively reassimilated fermentation-derived CO_2_ but also redirected carbon and reducing equivalents toward the synthesis of higher-value C6 products.

### 3.4. Enhanced carbon conservation and chain elongation through co-utilization of glucose, H_2_, and CO_2_ in fed-batch fermentation

As demonstrated in the batch fermentation, H_2_ supplementation enabled efficient carbon conservation and promoted carbon redistribution toward C6 products in ALECO2 (Fig. 2). To further evaluate the potential of ALECO2 for carbon conservation and chain elongation, fed-batch fermentations were performed under three different conditions. First, glucose-fed mixotrophic fermentation (F_mixo) was conducted by supplying 3 g/L glucose every 24 h, allowing internally generated CO_2_ to be reassimilated without external gas supplementation. This condition served as the control. Second, glucose/H_2_-fed mixotrophic fermentation (F_H_2__mixo) was performed by periodic supplementation of glucose (3 g/L) and H_2_ (150 kPa) to evaluate the effects of H_2_ supplementation on CO_2_ reassimilation and carbon flux redistribution toward C6 products under fed-batch conditions. Third, a glucose/CO_2_/H_2_-fed mixotrophic fermentation (F_CO_2_/H_2__mixo) was conducted to investigate whether the CO_2_-utilization capacity of ALECO2 extends beyond reassimilation of glucose-derived CO_2_ to the assimilation of externally supplied CO_2_. In this condition, glucose (3 g/L) and H_2_ (150 kPa) were supplied every 24 h, whereas CO_2_ was initially supplied at a partial pressure of 20 kPa and replenished to 20 kPa whenever a decrease in headspace CO_2_ pressure was observed. To mitigate product inhibition by C6 compounds and thereby sustain active chain elongation, GSP-25 resin was added for *in situ* adsorption of C6 compounds (Oh et al., 2026).

Under the F_mixo conditions, ALECO2 rapidly consumed each glucose pulse supplied during 144 h of fermentation and total glucose consumption was 20.80 g/L by 264 h (Fig. 3A). CO_2_ continuously accumulated in the headspace up to 50.6 kPa, whereas H_2_ increased only until approximately 72 h and subsequently remained relatively constant (Fig. 3A). These results suggest that CO_2_ generation from glucose metabolism exceeded its reassimilation capacity under F_mixo conditions. AA in the broth gradually increased to a maximum concentration of 3.43 g/L at 96 h and subsequently decreased to 2.80 g/L at the end of fermentation (Fig. 3D). After 96 h, no noticeable increase in metabolite concentrations was detected in the broth despite continued glucose consumption. This observation was likely attributable to adsorption of relatively hydrophobic fermentation products such as C4-C6 acids/alcohols onto the GSP-25 resin. When metabolites recovered from both the broth and resin were combined, AA was the predominant product with a final titer of 3.04 g/L, followed by HA at 2.75 g/L (Fig. 3G). Consistent with the continuous accumulation of CO_2_, the C_M_/C_S_ remained relatively low at 0.60, below the theoretical value of 0.67 expected from heterotrophic fermentation (Fig. 4D). This result indicates that carbon conservation through CO_2_ reassimilation was limited under F_mixo conditions.

**Figure 3.**
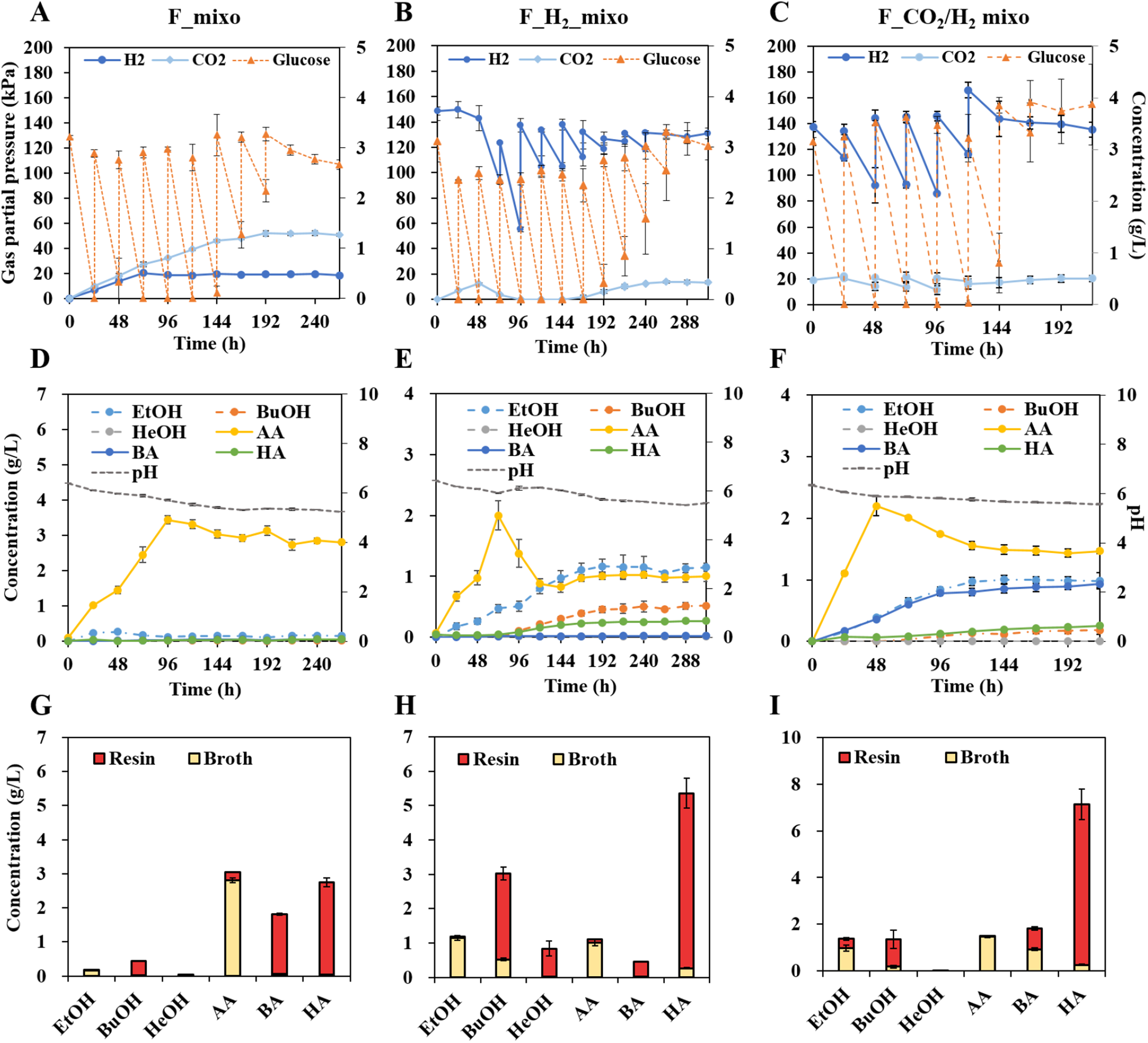
Fermentation performance of the ALECO2 strain under mixotrophic fed-batch conditions. (A-C) H_2_ and CO_2_ partial pressures and residual glucose concentrations during fed-batch fermentation of ALECO2 under (A) F_mixo, (B) F_H_2__mixo, and (C) F_CO_2_/H_2__mixo. (D-F) Time-course profiles of major fermentation products detected in the broth including ethanol (EtOH), butanol (BuOH), hexanol (HeOH), acetic acid (AA), butyric acid (BA), and hexanoic acid (HA), along with pH changes under each respective condition. (G-I) Final concentrations of metabolites detected in the broth and resin for each condition. Error bars represent the standard deviation of three biological replicates (n = 3).

**Figure 4.**
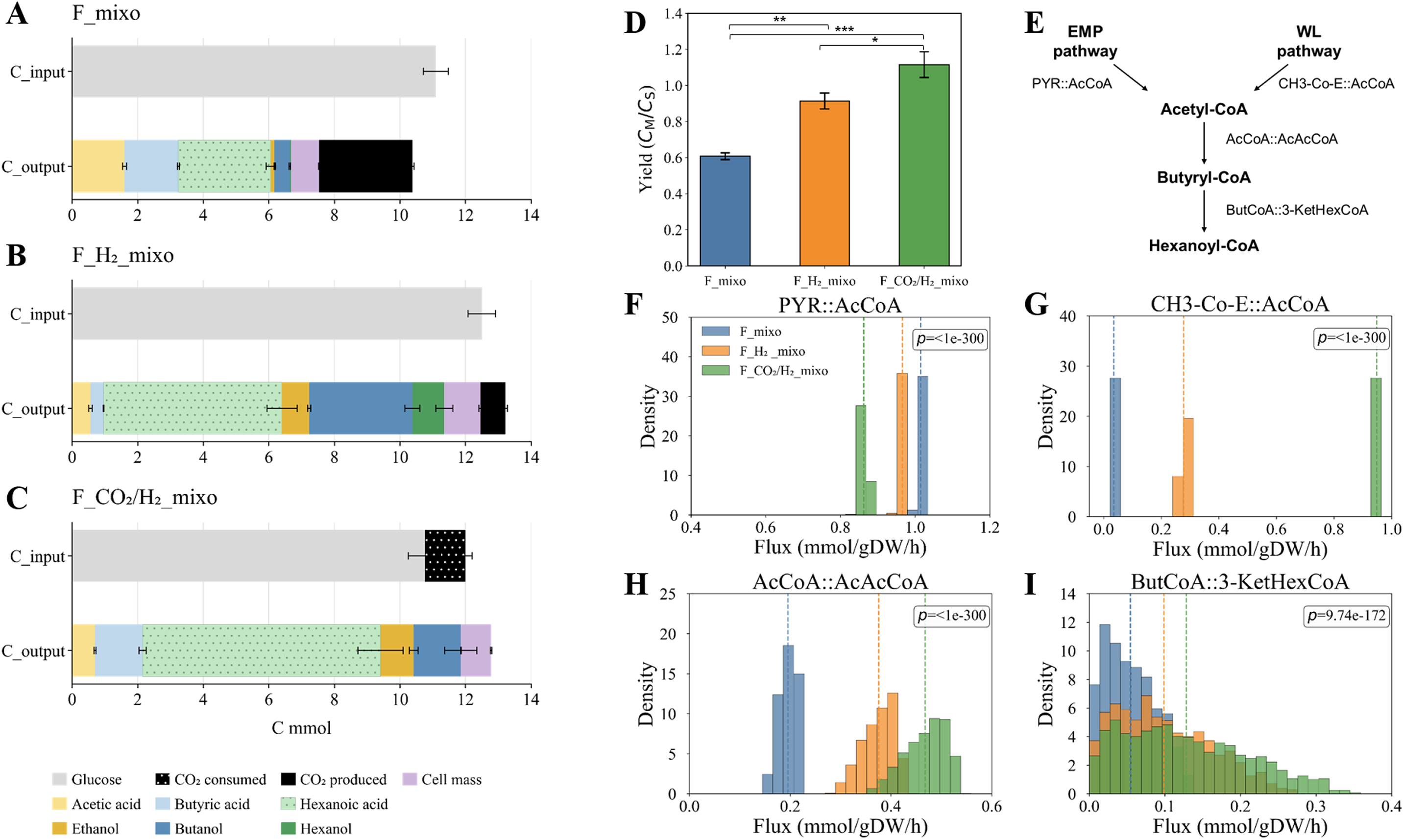
Carbon distribution, carbon molar yield (C_M_/C_S_), and intracellular flux distributions of the ALECO2 strain under different fed-batch fermentation conditions. (A-C) Carbon distribution in total C_input and total C_output during (A) F_mixo, (B) F_H_2__mixo, and (C) F_CO_2_/H_2__mixo. C_input was defined as the apparent carbon consumed from glucose and externally supplied CO_2_, whereas C_output included carbon recovered in fermentation products, biomass, and net CO_2_ produced in the headspace. (D) C_M_/C_S_ for each condition. Error bars indicate standard deviations of three biological replicates (n = 3). Asterisks indicate statistically significant differences among groups as determined by one-way ANOVA followed by Tukey’s post hoc test (\**p* < 0.05, \*\**p* < 0.01, \*\*\**p* < 0.001). (E) Schematic representation of the central metabolic pathways highlighting the positions of key acetyl-CoA-related reactions. (F-I) Histograms show the probability density distributions of sampled fluxes for key acetyl-CoA-related reactions (PYR::AcCoA, pyruvate-to-acetyl-CoA conversion; CH3-Co-E::AcCoA, acetyl-CoA formation via the Wood-Ljungdahl pathway; AcCoA::AcAcCoA and ButCoA::3-KetHexCoA, acetyl-CoA channeling into chain elongation). Dashed lines indicate mean flux values. Legends are shown only in panels A and F and apply to all corresponding panels. EMP, Embden-Meyerhof-Parnas pathway; WL, Wood-Ljungdahl pathway.

Under F_H_2__mixo conditions, glucose consumption and gas utilization continued up to 312 h, resulting in a total glucose consumption of 23.44 g/L (Fig. 3B). The final CO_2_ in the headspace was 13.4 kPa, markedly lower than that observed under F_mixo conditions (50.6 kPa), indicating that H_2_ supplementation enhanced CO_2_ reassimilation. Notably, CO_2_ accumulation was nearly absent between 96 and 144 h despite continued glucose consumption, suggesting that the rate of CO_2_ reassimilation exceeded the rate of CO_2_ generation from glucose metabolism during this period.

The product profile also differed markedly from that observed under F_mixo conditions. In the broth, AA accumulated until 72 h and subsequently decreased until 144 h, whereas EtOH, BuOH, HA, and HeOH continued to increase (Fig. 3E), suggesting redistribution of carbon from C2 compounds toward chain elongation and alcohol formation. When metabolites recovered from both the broth and resin were combined, HA became the predominant product with a final titer of 5.36 g/L (Fig. 3H), accounting for 41% (5.44 C-mmol) of total C_output (Fig. 4B). In F_mixo, C6 products accounted for only 28% of total C_output (2.86 C-mmol) (Fig. 4A), whereas the fraction increased to 49% (6.41 C-mmol) under F_H_2__mixo conditions (Fig. 4B), demonstrating a redistribution of carbon flux toward C6 product formation under F_H_2__mixo conditions. In addition, alcohols accounted for 37% of total C_output under F_H_2__mixo conditions compared with only 6% under F_mixo conditions, demonstrating that the additional reducing power supplied by H_2_ favored the formation of alcohols under F_H_2__mixo conditions. Consistent with the enhanced CO_2_ reassimilation, C_M_/C_S_ reached 0.91, substantially higher than that observed under F_mixo conditions (0.60) and approaching the theoretical maximum value of 1.0 (Fig. 4D). These results demonstrate that H_2_ supplementation not only effectively improved carbon conservation but also redirected carbon flux toward C6 compounds and alcohol production.

As shown in F_H_2__mixo conditions, CO_2_ accumulation was nearly absent between 96 and 144 h despite continued glucose consumption, suggesting that the rate of CO_2_ reassimilation exceeded the rate of CO_2_ generation from glucose metabolism (Fig. 3B). Based on this observation, additional CO_2_ was supplied under F_CO_2_/H_2__mixo conditions to determine whether ALECO2 could further utilize exogenous CO_2_ in addition to metabolically generated CO_2_, thereby increasing carbon recovery and potentially enabling carbon-negative fermentation. During the first 24 h, rapid glucose consumption was accompanied by an increase in headspace CO_2_ pressure (Fig. 3C). Subsequently, CO_2_ pressure continuously decreased and was repeatedly replenished to approximately 20 kPa. A consistent decline in CO_2_ pressure was observed following each replenishment cycle up to 120 h, suggesting active utilization of both glucose-derived CO_2_ and externally supplied CO_2_. After 120 h, CO_2_ consumption gradually slowed, resulting in a residual CO_2_ pressure of approximately 20 kPa at the end of fermentation. In total, 1.26 C-mmol of CO_2_ (22.5 kPa) was consumed together with glucose (20.16 g/L, 12.50 C-mmol) throughout the fermentation.

The product profile also shifted notably under F_CO_2_/H_2__mixo conditions. In the broth, AA accumulated until 48 h and subsequently decreased, whereas BA and EtOH continued to increase (Fig. 3F). When metabolites recovered from both the broth and resin were combined, HA was identified as the predominant product with a final titer of 7.14 g/L (Fig. 3I). The C_M_/C_S_ value reached 1.10, exceeding the theoretical maximum value of 1.0 expected from complete conservation of glucose-derived carbon alone (Fig. 4D). This result strongly suggests that, in addition to reassimilation of metabolically generated CO_2_, externally supplied CO_2_ was incorporated into fermentation products. Carbon distribution analysis further revealed enhanced chain elongation under F_CO_2_/H_2__mixo conditions. C6 products (7.26 C-mmol) accounted for 57% of total C_output (12.79 C-mmol), significantly higher than the 49% observed under F_H_2__mixo conditions (Welch’s t-test, *p* < 0.05). Overall, ALECO2 was capable of utilizing externally supplied CO_2_ in addition to glucose-derived CO_2_, resulting in net carbon uptake and carbon-negative fermentation during mixotrophic growth while maintaining active chain elongation.

### 3.5. In silico analysis of intracellular carbon flux redistribution

To assess whether the experimentally observed shifts in carbon distribution and product profiles were reflected at the intracellular metabolic level, constraint-based random sampling was performed using the core metabolic model of *C. carboxidivorans* P7 developed by Vees et al. (2022) with condition-specific constraints derived from fermentation data. The *C. carboxidivorans* P7 model was selected because this organism shares 99.9% 16S rRNA gene sequence similarity with JS66 and exhibits a similar fermentation phenotype, producing C2-C6 acids and alcohols (Kim et al., 2022).

Flux distributions were examined for reactions associated with acetyl-CoA generation and chain elongation, including PYR::AcCoA, representing glycolysis-derived acetyl-CoA formation; CH3-Co-E::AcCoA, representing WLP-derived acetyl-CoA formation; AcCoA::AcAcCoA, representing entry into reverse β-oxidation; and ButCoA::3-KetHexCoA, representing elongation toward C6 products (Fig. 4E). Condition-specific constraints were imposed using experimentally measured glucose uptake rates, H_2_ and CO_2_ exchange rates, and AA and EtOH production rates, all expressed in mmol/gDCW/h. CO_2_ exchange was constrained as production under F_mixo and F_H_2__mixo conditions and as consumption under F_CO_2_/H_2__mixo conditions. Because *in situ* resin adsorption limited accurate quantification of hydrophobic products, including C4-C6 acids and alcohols, throughout the fermentation, only AA and EtOH fluxes were used as product constraints. These values were calculated from the initial production phase, during which product accumulation remained linear. The growth rate was constrained to at least 90% of the condition-specific maximum growth rate obtained by FBA.

Under F_mixo conditions, acetyl-CoA formation was dominated by PYR::AcCoA, whereas CH3-Co-E::AcCoA contributed minimally, indicating limited WLP-derived acetyl-CoA formation (Fig. 4F-G). Fluxes through AcCoA::AcAcCoA and ButCoA::3-KetHexCoA were also relatively low (Fig. 4H-I). In contrast, flux distributions for CH3-Co-E::AcCoA were shifted toward higher values under the F_H_2__mixo and F_CO_2_/H_2__mixo conditions, indicating enhanced acetyl-CoA formation via the WLP (Kruskal-Wallis test, *p* < 0.001) (Fig. 4G). These conditions also exhibited increased sample flux distributions through AcCoA::AcAcCoA and ButCoA::3-KetHexCoA, suggesting that the additional WLP-derived acetyl-CoA was routed toward reverse β-oxidation and C6 chain elongation (Fig. 4H-I). This flux redistribution was most pronounced under F_CO_2_/H_2__mixo conditions and was consistent with the experimentally observed increases in C_M_/C_S_ (1.10) and C6 carbon fraction (57% of total C_output). Together, these model-based results qualitatively support that H_2_ and CO_2_/H_2_ supplementation enhanced WLP-derived acetyl-CoA formation and promoted carbon channeling toward chain-elongated C6 products.

Beyond enhanced carbon utilization, the flux redistribution observed under F_H_2__mixo and F_CO_2_/H_2__mixo conditions has important implications for medium-chain carboxylate production. The increased routing of carbon through WLP-derived acetyl-CoA formation and reverse β-oxidation suggests that these mixotrophic conditions can simultaneously improve carbon utilization and promote chain elongation toward higher-value C6 products. From a process perspective, this behavior is particularly attractive because it was achieved within a single microbial chassis. Previous bioproduction strategies based on cofeeding or co-culture systems often require fine-tuned substrate control or complex interspecies coordination (Charubin et al., 2019; Maru et al., 2018; Park et al., 2019). In contrast, the ALECO2 strain simultaneously supported CO_2_ assimilation and medium-chain acid biosynthesis under glucose-rich conditions without requiring microbial co-cultivation. This evolution-driven flexibility provides a promising platform for efficient mixotrophic carbon conversion coupled with the production of value-added medium-chain products.

### 3.6. Genomic basis of enhanced CO_2_ utilization in ALECO2

To identify the genetic basis underlying the enhanced CO_2_ utilization capacity of ALECO2, WGrS and *de novo* genome assembly were performed. The genome sequences of the ALECO70 and the adapted strain ALECO2 were compared against the deposited genome of JS66. In the WGrS analysis, seven mutations were identified in ALECO2 relative to ALECO70 (Table 1). One notable mutation was a 41-bp insertion in Q6H37_RS05210, which is annotated as encoding a NADH-ubiquinone oxidoreductase-F iron-sulfur binding region domain-containing protein, resulting in a frameshift and premature translation termination. The predicted Q6H37_RS05210 protein shares 86% amino acid sequence identity with HytB from both *C. autoethanogenum* and *C. ljungdahlii*, a subunit of the electron-bifurcating Hyt hydrogenase complex (HytABCDE_1_E_2_). Interestingly, mutations affecting Hyt hydrogenase complex components have been reported in acetogenic clostridia with enhanced H_2_/CO_2_ metabolism. For example, *hytA* deletion in *C. ljungdahlii* improved growth, H_2_ consumption, and AA and EtOH production under H_2_/CO_2_ conditions (Wen et al., 2026). Similarly, an evolved *C. carboxidivorans* strain selected for enhanced CO_2_ consumption carried a frameshift mutation in *hytA* that was proposed to impair the hexameric Hyt complex (Antonicelli et al., 2023). Although the ALECO2 mutation affects a putative *hytB* homolog rather than *hytA*, both genes encode components of the Hyt hydrogenase complex. The frameshift and premature termination in Q6H37_RS05210 may have structurally altered the encoded protein, affecting Hyt-dependent electron transfer during H_2_-dependent CO_2_ fixation, although its direct functional contribution remains to be experimentally validated. Two additional missense mutations were identified in a gene encoding a LysR family transcriptional regulator (LTTRs, Q6H37_RS15890). LTTRs are widely known as one of the most abundant families of transcriptional regulators in prokaryotes, involved in the regulation of diverse metabolic, stress response, and redox-related genes (Maddocks et al., 2008). Although the role of LTTRs has not been extensively characterized in *Clostridium* species, they have been reported to play important roles in CO_2_ fixation in autotrophic microorganisms such as *Rhodobacter capsulatus* and *Alcaligenes eutrophus* H16 (Van Keulen et al., 1998; Windhöver et al., 1991).

**Table 1.**
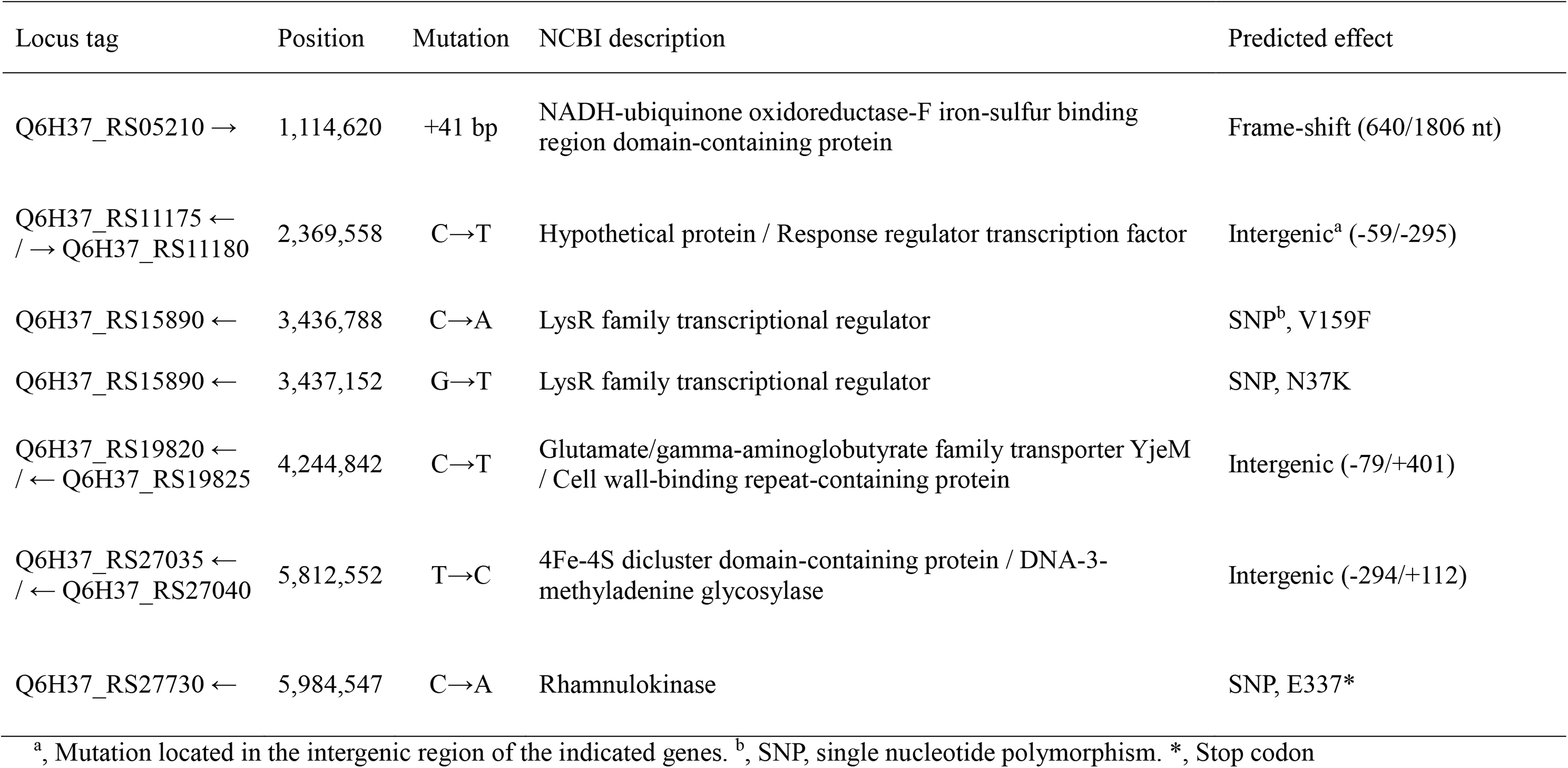
Mutations identified in the ALECO2 strain compared to ALECO70.

| Locus tag | Position | Mutation | NCBI description | Predicted effect |
| --- | --- | --- | --- | --- |
| Q6H37_RS05210 → | 1,114,620 | +41 bp | NADH-ubiquinone oxidoreductase-F iron-sulfur binding region domain-containing protein | Frame-shift (640/1806 nt) |
| Q6H37_RS11175 ←<br>/ → Q6H37_RS11180 | 2,369,558 | C→T | Hypothetical protein / Response regulator transcription factor | Intergenic<sup>a</sup> (-59/-295) |
| Q6H37_RS15890 ← | 3,436,788 | C→A | LysR family transcriptional regulator | SNP<sup>b</sup>, V159F |
| Q6H37_RS15890 ← | 3,437,152 | G→T | LysR family transcriptional regulator | SNP, N37K |
| Q6H37_RS19820 ←<br>/ ← Q6H37_RS19825 | 4,244,842 | C→T | Glutamate/gamma-aminobutyrate family transporter YjeM / Cell wall-binding repeat-containing protein | Intergenic (-79/+401) |
| Q6H37_RS27035 ←<br>/ ← Q6H37_RS27040 | 5,812,552 | T→C | 4Fe-4S dicluster domain-containing protein / DNA-3-methyladenine glycosylase | Intergenic (-294/+112) |
| Q6H37_RS27730 ← | 5,984,547 | C→A | Rhamnulokinase | SNP, E337* |

<sup>a</sup>, Mutation located in the intergenic region of the indicated genes. <sup>b</sup>, SNP, single nucleotide polymorphism. \*, Stop codon

Recent studies have shown that adaptive evolution can involve large-scale structural genome changes, including transposase-mediated genome restructuring during adaptation of *Thermoanaerobacter kivui* to CO metabolism (Hocq et al., 2025). Accordingly, structural genome variation was also examined between ALECO70 and ALECO2, and no large-scale structural changes were detected. Notably, comparison with JS66 revealed a chromosomal deletion of approximately 63.1 kb in both ALECO70 and ALECO2 (Fig. 5), suggesting that this deletion occurred during the earlier CO adaptation process rather than during the subsequent CO_2_/H_2_ adaptation. The deleted region included 72 predicted genes spanning positions 3,991,946-4,055,036 bp, including hypothetical proteins, transcriptional regulators, transport-related proteins, stress response proteins, structural RNA genes (5S, 16S, and 23S rRNA genes and a tRNA-Met gene), and cell wall-associated functions (Supplementary material-Excel file). Although the deleted region contained no genes directly involved in the WLP or hydrogenase complexes, the loss of stress response proteins and transcriptional regulators may have indirectly influenced the cellular environment or regulatory landscape in ways that facilitated adaptation to gas-utilizing growth, although the underlying mechanisms remain unclear.

**Figure 5.**
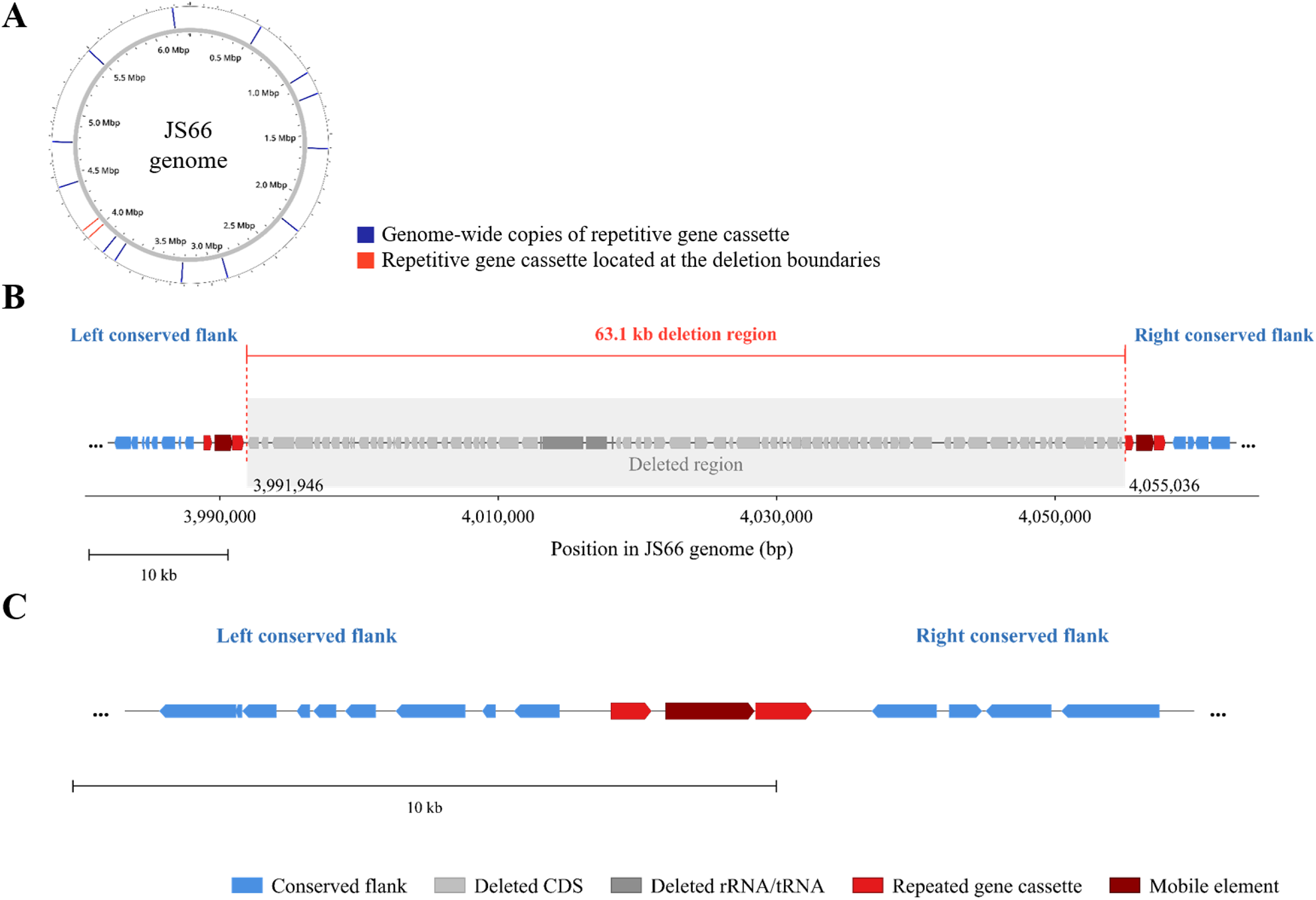
Genomic context of the 63.1 kb deletion in the evolved strains. (A) Circular map of the *Clostridium* sp. JS66 genome showing genome-wide repetitive gene cassettes and the repetitive gene cassettes located at the deletion boundaries. (B) Linear map of the 63.1 kb deletion region. Conserved flanking regions are shown in blue, deleted coding sequences and rRNA/tRNA genes in light and dark gray, respectively, repetitive gene cassettes in red, and the mobile element-associated gene in dark red. (C) Chromosomal structure of ALECO70 and ALECO2 after deletion.

Inspection of the deletion boundaries revealed that the deleted region was bordered by highly similar repetitive gene cassettes. Genome-wide analysis further showed that this cassette was present in 15 copies across the JS66 chromosome (Fig. 5A and Supplementary material-Excel file). These repetitive gene cassettes shared a conserved three-gene organization, consisting of genes encoding a DUF6431 domain-containing protein, a DDE (Asp-Asp-Glu catalytic motif)-type integrase/transposase/recombinase, and an AAA family ATPase (ATPases associated with diverse cellular activities) (Fig. 5B and Supplementary material-Excel file). In the genome of ALECO70 and ALECO2, the two repetitive gene cassettes bordering the deleted segment appeared to have been resolved into a single retained copy (Fig. 5C), resembling the structural outcome commonly observed in homologous recombination-mediated deletions (Reams et al., 2012).

Collectively, the genomic alterations identified in ALECO2 suggest that enhanced CO_2_ utilization may have resulted from multiple layers of adaptation, including mutations related to electron transfer and transcriptional regulation. Future studies combining targeted genetic manipulation with transcriptomic and physiological characterization will be necessary to determine the functional contribution of each mutation and to elucidate the molecular mechanisms underlying the evolved CO_2_ utilization phenotype.

## 4. Conclusion

This study established ALECO2 as an evolved anaerobic non-photosynthetic (ANP) mixotrophy platform that couples glucose fermentation with H_2_-assisted CO_2_ reassimilation and chain elongation. Under H_2_-mixotrophic batch conditions, ALECO2 nearly completely reassimilated glucose-derived CO_2_, achieving a carbon molar yield (C_M_/C_S_) of 0.95. Under fed-batch conditions, H_2_ supplementation increased C_M_/C_S_ from 0.60 to 0.91 while increasing the C6 carbon fraction from 28% to 49% compared with glucose-fed fermentation. Supplementation with both H_2_ and CO_2_ further increased C_M_/C_S_ to 1.10, demonstrating carbon-negative fermentation while increasing the C6 carbon fraction to 57%. Genome analysis identified mutations in genes encoding a putative HytB homolog and a LysR-type transcriptional regulator. To the best of our knowledge, this is the first report describing an evolved acetogenic monoculture that integrates ANP mixotrophy, external CO_2_ assimilation, and enhanced C6 production. These findings highlight ALECO2 as a promising platform for carbon-negative production of value-added medium-chain products.

## Supporting information

Supplementary materials_Word

## Data availability

Data will be made available on request.

## E-supplementary data

E-supplementary data can be found in e-version of this paper online.

## Declaration of generative AI and AI-assisted technologies in the manuscript preparation process

During the preparation of this work, the authors used ChatGPT (OpenAI) in order to improve the English grammar, clarity and conciseness of written text. After using this tool, the authors reviewed and edited the content as needed and take full responsibility for the content of the published article.

## References

Antonicelli, G., Ricci, L., Tarraran, L., Fraterrigo Garofalo, S., Re, A., Vasile, N.S., Verga, F., Pirri, C.F., Menin, B., Agostino, V., 2023. Expanding the product portfolio of carbon dioxide and hydrogen-based gas fermentation with an evolved strain of *Clostridium carboxidivorans*. Bioresour. Technol. 387, 129689.

Braun, M., Mayer, F., Gottschalk, G., 1981. *Clostridium aceticum* (Wieringa), a microorganism producing acetic acid from molecular hydrogen and carbon dioxide. Arch. Microbiol. 128(3), 288–93.

Charubin, K., Papoutsakis, E.T., 2019. Direct cell-to-cell exchange of matter in a synthetic *Clostridium* syntrophy enables CO_2_ fixation, superior metabolite yields, and an expanded metabolic space. Metab. Eng. 52, 9–19.

Chin, C.-S., Alexander, D.H., Marks, P., Klammer, A.A., Drake, J., Heiner, C., Clum, A., Copeland, A., Huddleston, J., Eichler, E.E., Turner, S.W., Korlach, J., 2013. Nonhybrid, finished microbial genome assemblies from long-read SMRT sequencing data. Nat. Methods. 10(6), 563–569.

Deatherage, D.E., Barrick, J.E. 2014. Identification of mutations in laboratory-evolved microbes from next-generation sequencing data using breseq. in: Engineering and analyzing multicellular systems: methods and protocols, Springer, pp. 165–188.

Deng, Z., Zhu, B., Davis, S.J., Ciais, P., Liu, Z., 2026. Global carbon emissions and decarbonization in 2025. Nat. Rev. Earth Environ. 7(5), 274–276.

Ebrahim, A., Lerman, J.A., Palsson, B.O., Hyduke, D.R., 2013. COBRApy: COnstraints-Based Reconstruction and Analysis for Python. BMC Syst. Biol. 7(1), 74.

Fagan, R.P., Fairweather, N.F., 2011. *Clostridium difficile* has two parallel and essential Sec secretion systems. J. Biol. Chem. 286(31), 27483–27493.

Fast, A.G., Schmidt, E.D., Jones, S.W., Tracy, B.P., 2015. Acetogenic mixotrophy: novel options for yield improvement in biofuels and biochemicals production. Curr. Opin. Biotechnol. 33, 60–72.

Hocq, R., Horvath, J., Stumptner, M., Malevičius, M., Thallinger, G.G., Pflügl, S., 2025. A megatransposon drives the adaptation of *Thermoanaerobacter kivui* to carbon monoxide. Nat. Commun. 16(1), 4217.

Jones, S.W., Fast, A.G., Carlson, E.D., Wiedel, C.A., Au, J., Antoniewicz, M.R., Papoutsakis, E.T., Tracy, B.P., 2016. CO_2_ fixation by anaerobic non-photosynthetic mixotrophy for improved carbon conversion. Nat. Commun. 7(1), 12800.

Jung, K.H., Lee, J., Lee, J., Ahn, J.H., Ko, J.K., Gong, G., Lee, S.-M., Han, S.O., Um, Y., 2026. Hexanoic acid production from lignocellulosic hydrolysates via simultaneous glucose–xylose utilization by carbon catabolite repression-relaxed *Clostridium* sp. JS66 with *in situ* adsorption. Bioresour. Technol. 457, 134986.

Kim, J., Kim, K.-Y., Ko, J.K., Lee, S.-M., Gong, G., Kim, K.H., Um, Y., 2022. Characterization of a novel acetogen *Clostridium* sp. JS66 for production of acids and alcohols: focusing on hexanoic acid production from syngas. Biotechnol. Bioprocess Eng. 27(1), 89–98.

Kolmogorov, M., Yuan, J., Lin, Y., Pevzner, P.A., 2019. Assembly of long, error-prone reads using repeat graphs. Nat. Biotechnol. 37(5), 540–546.

Li, W., Cheng, C., Zhao, J., Li, Q., Liu, M., Wang, C., Xue, C., 2024. Enhanced carbon recovery and butyrate production in cocultivation of *Clostridium tyrobutyricum* and *C. ljungdahlii*. ACS Sustain. Chem. Eng. 12(35), 13244–13252.

Maddocks, S.E., Oyston, P.C., 2008. Structure and function of the LysR-type transcriptional regulator (LTTR) family proteins. Microbiology. 154(12), 3609–3623.

Maru, B.T., Munasinghe, P.C., Gilary, H., Jones, S.W., Tracy, B.P., 2018. Fixation of CO_2_ and CO on a diverse range of carbohydrates using anaerobic, non-photosynthetic mixotrophy. FEMS Microbiol. Lett. 365(8), fny039.

Megchelenbrink, W., Huynen, M., Marchiori, E., 2014. optGpSampler: an improved tool for uniformly sampling the solution-space of genome-scale metabolic networks. PLoS One. 9(2), e86587.

Oh, H.J., Ahn, J.H., Gong, G., Ko, J.K., Lee, S.-M., Um, Y., 2026. High hexanol production from syngas by *Clostridium carboxidivorans* P7 through *in situ* hexanol recovery by adsorption. Renew. Energy. 261, 125312.

Park, J.O., Liu, N., Holinski, K.M., Emerson, D.F., Qiao, K., Woolston, B.M., Xu, J., Lazar, Z., Islam, M.A., Vidoudez, C., 2019. Synergistic substrate cofeeding stimulates reductive metabolism. Nat. Metab. 1(6), 643–651.

Reams, A.B., Kofoid, E., Kugelberg, E., Roth, J.R., 2012. Multiple pathways of duplication formation with and without recombination (RecA) in *Salmonella enterica*. Genetics. 192(2), 397–415.

Song, X., Ju, Y., Chen, L., Zhang, W., 2024. Strategies and tools to construct stable and efficient artificial coculture systems as biosynthetic platforms for biomass conversion. Biotechnol. Biofuels Bioprod. 17(1), 148.

Van Keulen, G., Girbal, L., Van Den Bergh, E., Dijkhuizen, L., Meijer, W., 1998. The LysR-type transcriptional regulator CbbR controlling autotrophic CO_2_ fixation by *Xanthobacter flavus* is an NADPH sensor. J. Bacteriol. 180(6), 1411–1417.

Vees, C.A., Herwig, C., Pflügl, S., 2022. Mixotrophic co-utilization of glucose and carbon monoxide boosts ethanol and butanol productivity of continuous *Clostridium carboxidivorans* cultures. Bioresour. Technol. 353, 127138.

Wen, Z.-Q., Li, Y.-Z., Su, H.-J., Zhang, J.-Z., Liu, Z.-Y., Wang, S.-N., Wang, H., Li, F.-L., Ma, X.-Q., 2026. Deletion of electron-bifurcating [FeFe]-hydrogenase enhances H_2_-driven CO_2_ fixation and ethanol production in *Clostridium ljungdahlii*. Bioresour. Technol. 453, 134636.

Wijaya, Y.P., Putra, R.D.D., Widyaya, V.T., Ha, J.-M., Suh, D.J., Kim, C.S., 2014. Comparative study on two-step concentrated acid hydrolysis for the extraction of sugars from lignocellulosic biomass. Bioresour. Technol. 164, 221–231.

Willis, N.B., Otten, J.K., Seo, H., Munasinghe, P.C., Hill, J.D., Papoutsakis, E.T., 2026. Enabling supratheoretical isopropanol yields from carbon-negative glucose fermentations with a *Clostridium acetobutylicum*-*Clostridium ljungdahlii* coculture. Metab. Eng. 96, 92–103.

Windhöver, U., Bowien, B., 1991. Identification of *cfxR*, an activator gene of autotrophic CO_2_ fixation in *Alcaligenes eutrophus*. Mol. Microbiol. 5(11), 2695–2705.

