## Supplementary materials_Word for "Adaptive Laboratory Evolution (ALE) enables carbon-negative mixotrophic fermentation and enhanced chain elongation in *Clostridium* sp. JS66"

**Figure S1 for section 3.1**

**Figure S2 for section 3.1, 3.2**

**Figure S3 for section 3.2, 3.3**

**Figure S1 for section 3.1**

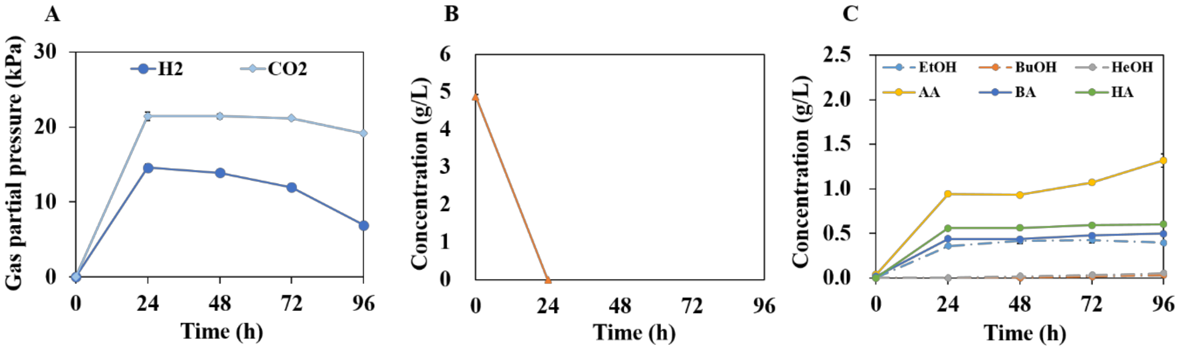


Figure S1. Fermentation profiles of *Clostridium* sp. JS66 with 5 g/L glucose. (A) Time-course profiles of $H_2$ and $CO_2$ production, (B) glucose consumption, and (C) Ethanol (EtOH), butanol (BuOH), hexanol (HeOH), acetic acid (AA), butyric acid (BA), and hexanoic acid (HA) production. Error bars represent the range of two biological replicates (n = 2).

**Figure S2 for section 3.1, 3.2**

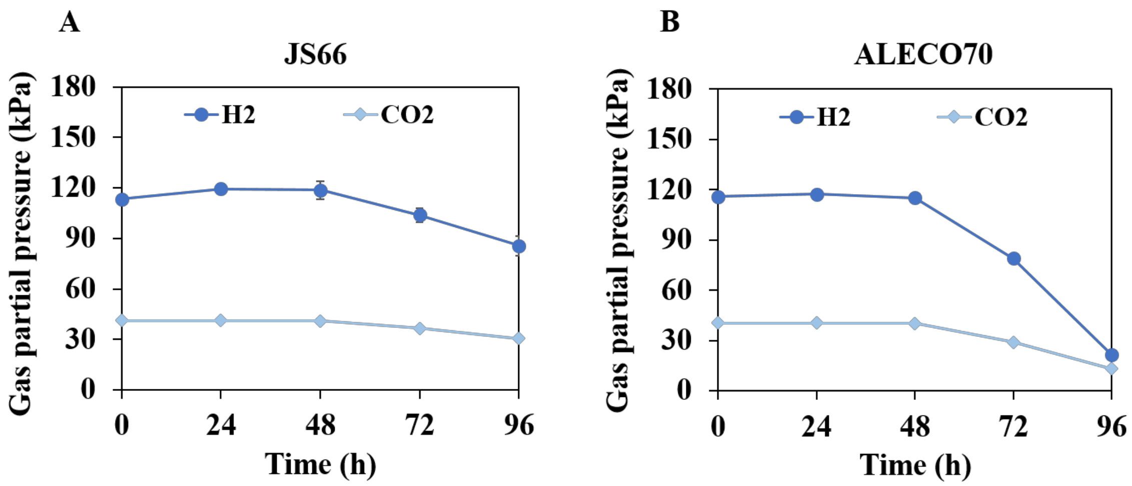


**Figure S2.** Gas utilization profiles of *Clostridium* sp. JS66 (JS66) and ALECO70 during $CO_2/H_2$ fermentation. Time-course profiles of headspace $H_2$ and $CO_2$ consumption by (A) JS66 and (B) ALECO70. Error bars represent the range of two biological replicates (n = 2).

**Figure S3 for section 3.2, 3.3**

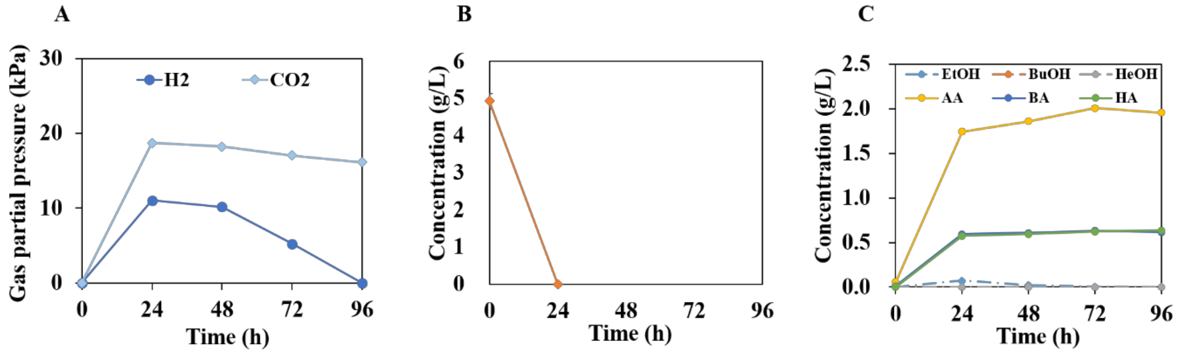


Figure S3. Fermentation profiles of ALECO2 with 5 g/L glucose. (A) Time-course profiles of $H_2$ and $CO_2$ production, (B) glucose consumption, and (C) Ethanol (EtOH), butanol (BuOH), hexanol (HeOH), acetic acid (AA), butyric acid (BA), and hexanoic acid (HA) production. Error bars represent the range of two biological replicates (n = 2).
